# Molecular measurable residual disease testing of blood during AML cytotoxic therapy for early prediction of clinical response

**DOI:** 10.1101/422352

**Authors:** Hong Yuen Wong, Anthony D. Sung, Katherine E. Lindblad, Sheenu Sheela, Gregory W. Roloff, David Rizzieri, Meghali Goswami, Matthew P. Mulé, Nestor R. Ramos, Jingrong Tang, Julie Thompson, Christin B. DeStefano, Kristi Romero, Laura W. Dillon, Dong-Yun Kim, Catherine Lai, Christopher S. Hourigan

## Abstract

**PURPOSE:** Measurable residual disease (MRD) testing after initial chemotherapy treatment can predict relapse and survival in AML. However, it has not been established if repeat molecular or genetic testing during chemotherapy can offer information regarding the chemotherapy sensitivity of the leukemic clone.

**PATIENTS AND METHODS:** Blood from 45 adult AML patients at day 1 and 4 of induction (n = 35) or salvage (n = 10) cytotoxic chemotherapy was collected for both quantitative real-time PCR (qPCR) assessment (*WT1*) and next generation sequencing (NGS, >500x depth) of 49 gene regions recurrently mutated in MDS/AML.

**RESULTS:** The median age was 62 (23-78); 42% achieved a complete response. *WT1* was overexpressed in most patients tested but was uninformative for very early MRD assessment. A median of 4 non-synonymous variants (range 0-7) were detected by DNA sequencing of blood on day 1 of therapy (median VAF: 29%). Only two patients had no variants detectable. All mutations remained detectable in blood on day 4 of intensive chemotherapy and remarkably the ratio of mutated to wild-type sequence was often maintained. This phenomenon was not limited to variants in *DNMT3A, TET2* and *ASXL1*. The kinetics of *NPM1* and *TP53* variant burden early during chemotherapy appeared to be exceptions and exhibited consistent trends in this cohort.

**CONCLUSIONS:** Molecular testing of blood on day 4 of chemotherapy is not predictive of clinical response in AML. The observed stability in variant allele frequency suggests that cytotoxic therapy may have a limited therapeutic index for clones circulating in blood containing these mutations. Further validation is required to confirm the utility of monitoring *NPM1* and *TP53* kinetics in blood during cytotoxic therapy.

## Introduction

The use of high sensitivity techniques to measure residual leukemic burden in patients achieving a complete remission by cytomorphological criteria is increasingly considered part of the standard of care for acute myeloid leukemia (AML) ^1-4^. While testing for measurable residual disease (MRD) in AML is typically performed using multi-parameter flow cytometry (MPFC) or real-time quantitative PCR (qPCR) there is increasing recent research interest in the potential of sequencing-based approaches^5-12^. The results of MRD testing in AML appear prognostic when measured at key landmark timepoints following initial therapy, typically after 1-2 cycles of induction therapy or before allogeneic transplantation^13-19^. It is currently not known if testing changes in residual leukemic burden at earlier timepoints, for example during initial induction therapy, would have clinical utility.

We therefore used two independent molecular techniques for AML MRD quantification, *WT1* expression by qPCR and targeted DNA sequencing for common MDS/AML variants, to determine if early assessment of blood from AML patients during the first four days of intensive cytotoxic therapy can predict subsequent clinical response.

## Methods

### Patients and sample collection

Blood was collected daily from day 1 through at least day 5 of intensive cytotoxic therapy from 45 adult AML patients with a median age of 62 years-old (range: 23-78) following informed consent on IRB-approved protocols (Table 1). Ten patients (median age: 52, range: 23-66) had relapsed or refractory AML (RR-AML) and were recruited to the National Heart, Lung, and Blood Institute (NHLBI) at the National Institutes of Health to receive salvage chemotherapy (NCT02527447). Thirty-five patients (median age: 63, range: 30-78) were treated at Duke University School of Medicine with induction chemotherapy for newly diagnosed AML (Figure S1). In addition, research samples from clinically indicated bone marrow examinations and post-induction follow-up visits were available for a subset of 22 patients.

**Table 1.**
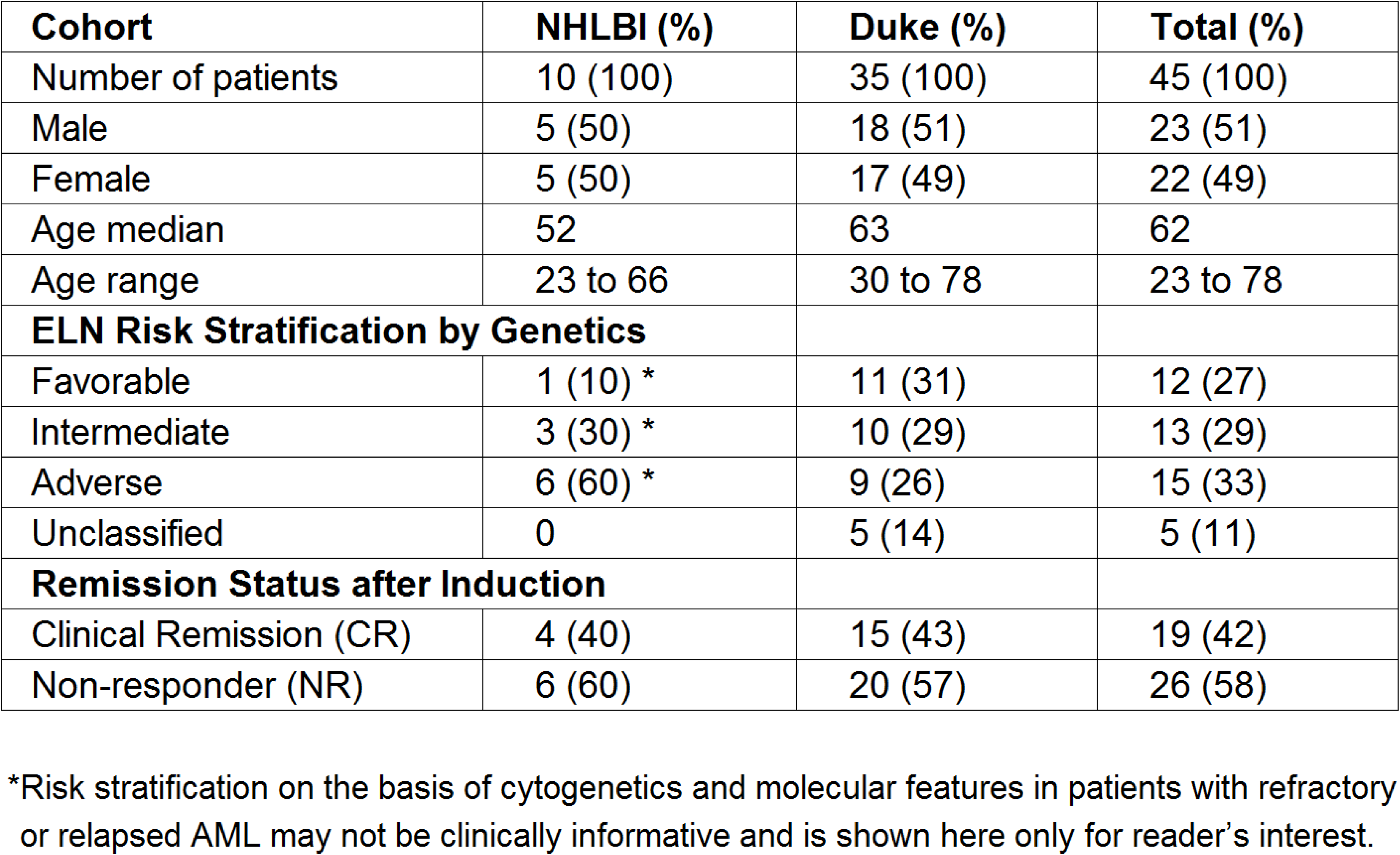
Patient Characteristics

### Nucleic acid extraction from blood and marrow

Samples were processed with NucleoSpin Blood QuickPure kits (Machery-Nagel) and NucleoSpin RNA Blood kits (Machery-Nagel), as per manufacturer’s instructions (NIH) or stored in PAXgene Blood/Marrow RNA tubes (PreAnalytiX) at Duke and shipped frozen to the NIH. Upon thawing PAXgene tubes, 2mL for gDNA isolation were pelleted and resuspended in PBS then processed with the QIAamp DNA blood mini kit (QIAGEN), as per manufacturer’s instructions. The remaining volume was processed with the PAXgene Blood RNA Kit (PreAnalytiX), per manufacturer’s instructions.

### Real-time quantitative PCR (qPCR)

RNA (130-260ng) was reverse transcribed using the RT^2^ First Strand kit (330404, QIAGEN). When necessary, RNA was concentrated with a Savant SVC-100H centrifugal evaporator. Resultant cDNA was loaded into a Custom RT^2^ Profiler PCR Array containing lyophilized qPCR primers for *WT1* and *ABL1* using the QIAgility System (QIAGEN). qPCR was performed (hold 2m at 50°C, hold 10m at 95°C, then 50 cycles of 15s at 95°C and 60s at 60°C) on the Rotor-Gene Q Platform (QIAGEN) and Ct values were collected with a threshold of 0.06. Healthy levels of *WT1* expression were established based on upper limit observed in blood of 34 healthy donors.

### DNA sequencing using a myeloid panel during chemotherapy

A total of 49 gene or gene regions recurrently mutated in MDS and/or AML were sequenced using amplicon-based targeted DNA sequencing (RainDance, Billerica, MA). Libraries were prepared from 100ng of gDNA and paired-end 300bp sequencing was performed on the MiSeq instrument (Ilumina), as per manufacturer’s instructions. Results were analyzed using the NextGENe v2.4.2.1 software (SoftGenetics, PA). Sequences were aligned to human genome build v37 (hg19). Noncoding and synonymous variants, along with known sequencing artifacts and regions with less than 500-fold coverage were removed. Remaining variants (i.e. missense, in-frame, frameshift, and nonsense mutations) with variant allele frequency (VAFs) above 5% at either day 1 or 4 were considered in subsequent analyses.

### Custom DNA sequencing for MRD tracking after chemotherapy

In order to track variants in longitudinal blood samples in 2 patients, a custom DNA sequencing assay was designed (DB0188, VariantPlex, ArcherDX). Libraries were prepared from 400ng gDNA; using paired-end 150bp sequencing on MiSeq instrument (Ilumina), as per manufacturer’s instructions. Archer Analysis software version 5.1.3 was used for analysis.

### Statistical analysis

All statistical analyses were performed using Prism v7.02 (GraphPad Software, CA). The Wilcoxson signed-rank test was performed on paired D1/D4 VAFs. Additive (D4 – D1, with denominator of D1 or D4) and multiplicative (D4/D1) differences between D1 and D4 were calculated, then the Wilcoxson rank-sum test was performed on these differences between CR/NR groups. The Chi-square test was performed based on whether the additive difference was positive or negative between CR and NR groups. The unpaired t-test was performed on relative expression levels of *WT1*. For all statistical tests, P<0.05 was considered significant.

## Results

### Patients and sample collection

Full demographics, risk classification, treatment and responses are listed in Tables 1, S1 and S2. Overall, 19 of 45 patients achieved a complete remission (CR) after intensive therapy (42%). Average white blood cell count was 10K/ul (range: 0.3-60) on day 1 decreasing to 2.3K/ul (range: 0.2-26) by treatment day 4. Obtaining sufficient quantities of nucleic acid from blood is the limiting factor for molecular testing during these early time points of cytotoxic chemotherapy in AML. All 45 patients had sufficient DNA for sequencing but only 34/45 patients had enough RNA from paired day 1 and 4 samples for qPCR analysis. There was insufficient RNA and DNA yields from blood beyond day 4 in most patients (Fig. S2).

### *WT1* expression level on day 4 is an uninformative biomarker of clinical response

*WT1* gene expression (normalized to *ABL1* expression) was determined in 34 patients with sufficient RNA for qPCR analysis isolated from blood on days 1 and 4 of treatment. Compared with the upper limit of expression observed in healthy donors (Figure 1A) 31 of 34 patients overexpressed *WT1* on Day 1 (91%) consistent with prior reports^20,21^. By day 4 of induction therapy 23 patients had *WT1* over-expression. 7 of 13 (54%) patients achieving a CR had at least a 4-fold reduction in *WT1* expression, although notably 6 of these 7 patients remained overexpressed compared with healthy donors on day 4 (Figure 1B). 7 of 21 (33%) patients who were non-responders (NR) also had at least a 4-fold reduction in *WT1* expression. Two NR patients with undetectable *WT1* on day 1 had low level expression on day 4 (Figure 1C). Overall, changes in *WT1* expression in blood between day 1 and 4 of intensive cytotoxic chemotherapy for AML appear uninformative as a biomarker for clinical response in this cohort.

**Figure 1.**
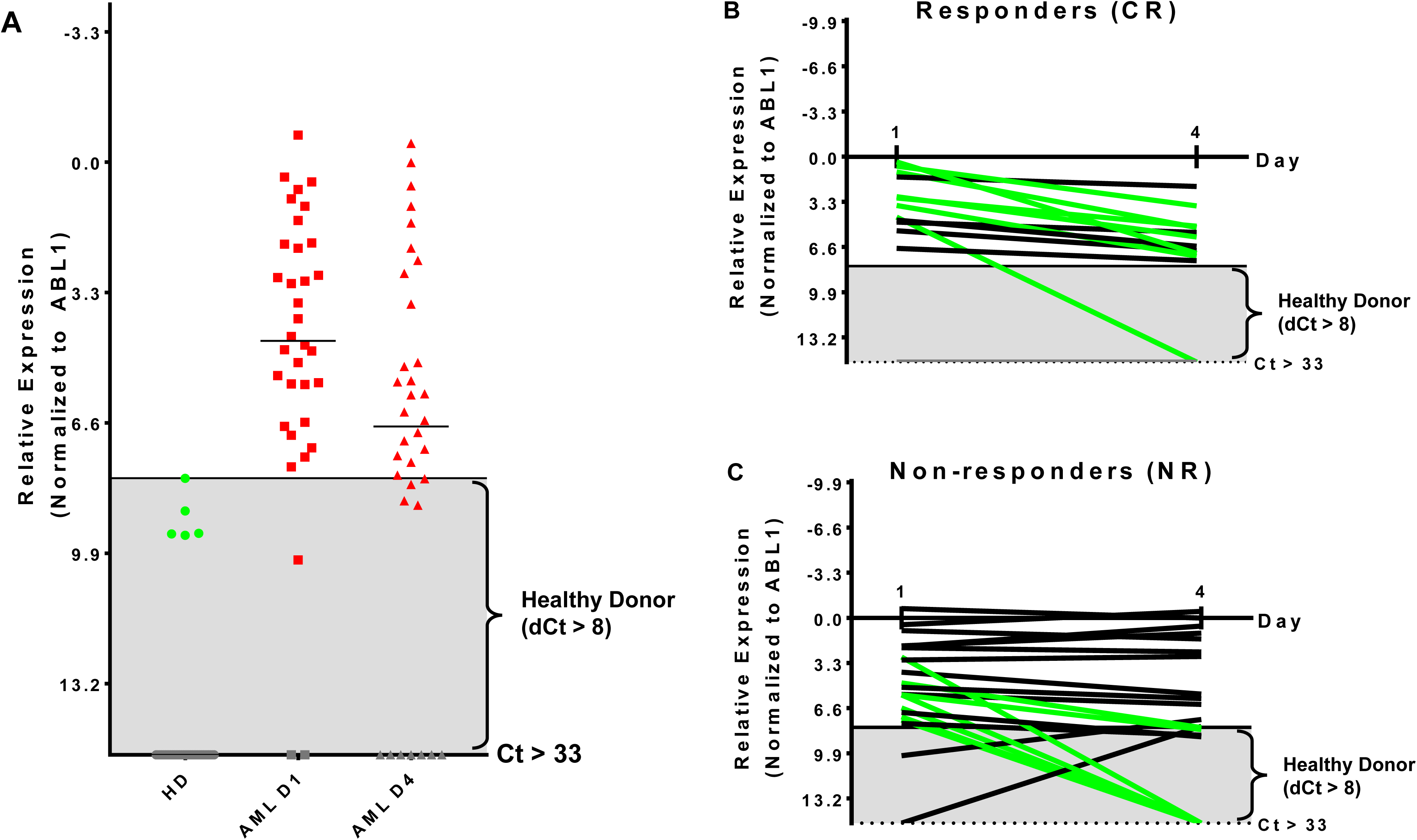
Kinetics of *WT1* expression during chemotherapy is uninformative for clinical response. **(A)** Thresholds for *WT1* overexpression in blood for this assay were established based on the upper limit observed by qPCR in 34 healthy donors and were consistent with previous reports^40^. AML patient blood samples showed overexpressed *WT1* levels in 31 (91%) on Day 1 and 23 (68%) at Day 4. **(B)** Patients achieving a complete remission after therapy had at least a 4-fold reduction of *WT1* expression during therapy in 7 of 13 cases evaluated by qPCR (54%); 5/13 (38%) had less than 4-fold reduction; 1/13 (8%) had undetectable *WT1* levels. **(C)** Non-responding patients had at least a 4-fold reduction of *WT1* expression in 7 of 21 evaluated by qPCR (33%) NR; 14/21 (67%) had less than a 4-fold change. Two patients initially had *WT1* levels that were not overexpressed, but became so by treatment day 4. **(D)** qPCR, quantitative real-time PCR; Green, at least 4-fold decrease; Black, less than 4-fold change; Gray, undetectable; Gray box indicates healthy donor range.

### DNA sequencing for variants associated with myeloid malignancy pre-treatment

All patients had targeted DNA sequencing (Table S3) of blood taken on day 1 and day 4 of chemotherapy. An average of 4 coding variants (range 0-7) were identified per patient, and only two patients had no variants suitable for disease tracking available. A total of 163 variants were found in 43 patients and some patients had multiple variants found within a single gene (considering multiple variants in one gene in the same patient as single event results in a total of 140 mutated genes). The most frequently mutated gene regions were consistent with those previously reported^22,23^ (see Figure S3AB). There was no difference in the number of coding variants detectable at baseline in CR vs. NR patients (Figure 2). Variant allele frequencies (VAF) on day 1 were a median of 29% (range: 0.2%-71%) with 75% having a VAF <40% (Figure S3C) on day 1. Consistent with current prognostic risk classifications^1^, the *TP53* and *NPM1* mutation classes had predictive significance, with 9 of 9 patients with *TP53* mutations not achieving remission while all 5 of 5 patients with *NPM1* mutations achieved CR.

**Figure 2.**
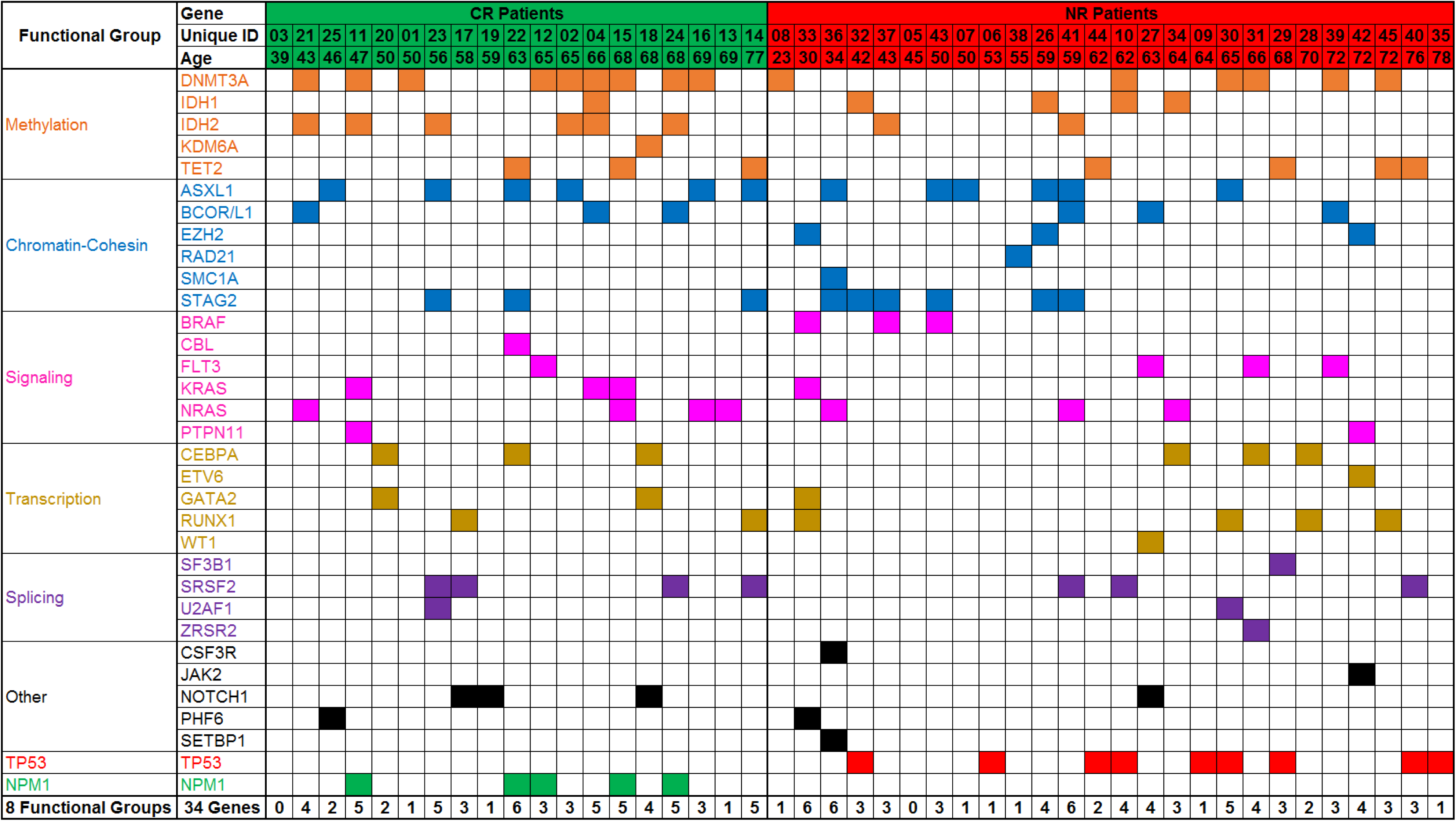
Targeted DNA sequencing of AML patient blood from day 1 of treatment. Variants were detected in 43 of 45 patients assessed. A total of 162 variants were identified in 34 genes or gene regions of 49 assessed. Genes with variants are grouped by gene function/class. Frequency of mutations and patterns of co-mutation are consistent with previous reports. CR: complete remission, NR: non-responder.

### Detectable variant allele frequency kinetics in blood during chemotherapy

Of the 43 patients with at least 1 variant detected in blood at day 1, 18 (42%) achieved CR and 25 (58%) were NR after induction therapy. Although white blood cell (WBC) count decreased 75% on average in the first 4 days of therapy (Figure S2A), all variants detected in the blood on day 1 of chemotherapy remained detectable in blood on day 4. VAFs on day 1 and day 4 were compared based on the hypothesis that changes in detectable mutation burden in blood very early during intensive cytotoxic treatment may correlate with clinical response as later assessed by morphological examination of bone marrow at count recovery (ie: approximately 30 days later). Day 1 and day 4 VAFs, at both the genetic functional class level and individual gene level, were compared using the Wilcoxon Sign Test, for all patients and also for the subgroups achieving either CR or NR (Figures 3 and S4AB). TP53 mutations were only detected in NR patients and showed a significant difference in VAF between days 1 and 4 (P<0.05) with a mean increase of 34% (n=9; range: −2% to 94%). NPM1 mutations were observed only in CR patients with a mean decrease of −44% (n=5; range: −2% to −98%) (P=0.0625). Furthermore, additional testing for 2-sample statistical significance of the additive and multiplicative differences was assessed between Day 1 and 4 VAFs at the genetic functional class level between the CR and NR patient groups, all of which were nonsignificant.

**Figure 3.**
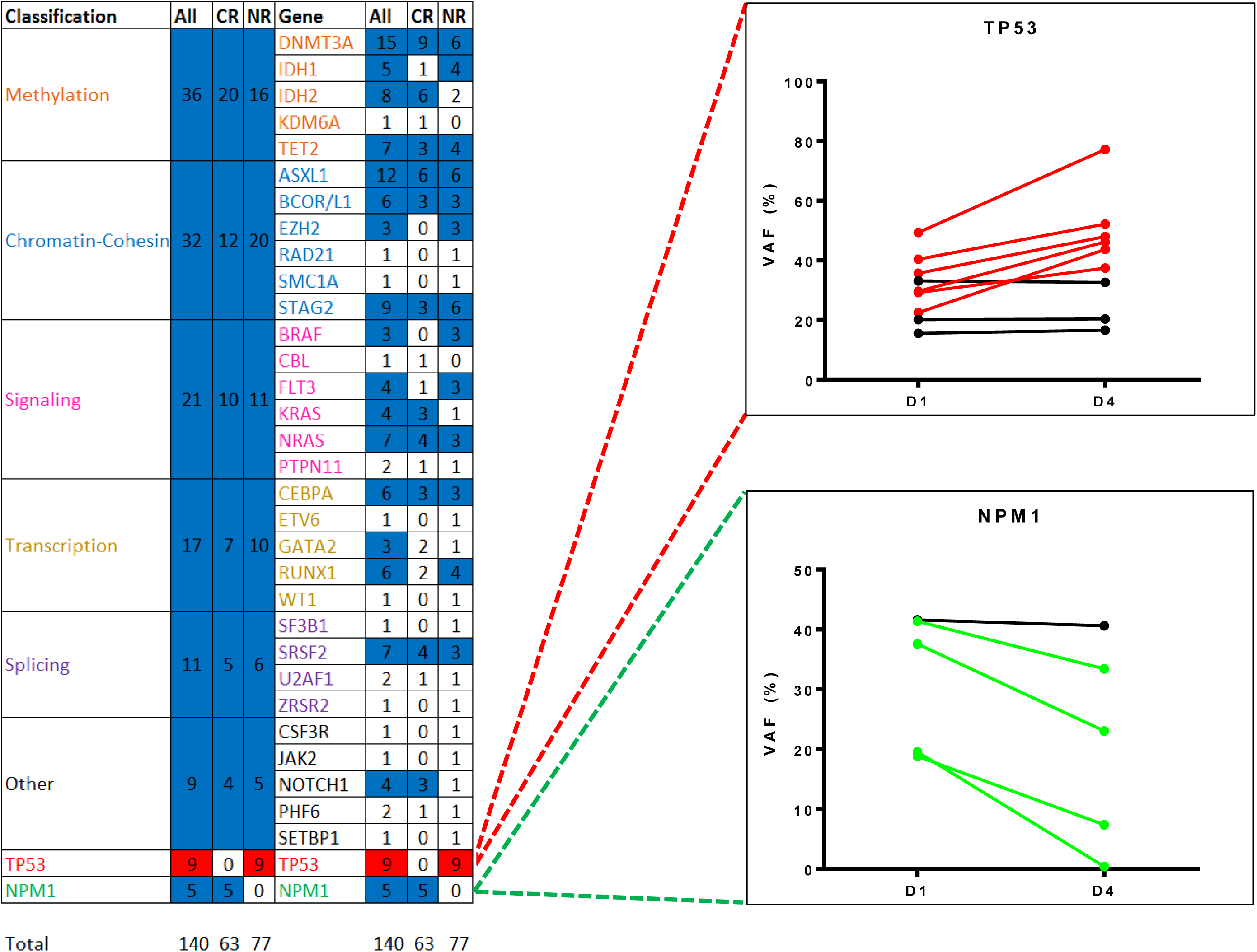
Changes in variant frequency during chemotherapy. All variants identified by targeted DNA sequencing in blood from day 1 of treatment (median VAF of 29%, range: 0.2%-71%) were also detectable in blood from day 4 (median VAF 27%, range: 0.1%-86%). Heatmap shows statistically significant changes in VAF between day 1 and day 4 by either gene functional class or by individual gene/gene region, for all patients or just those with complete remission (CR) or non-responder (NR). Variants were only counted once per gene region per patients. Mutated *TP53*, detected only in NR patients, was significantly different (red, P<0.05) and mutated *NPM1*, detected only in CR patients, demonstrated a consistent trend (blue, P=0.0625). The remaining functional groups and individual genes were either nonsignificant (blue) or had too few data points for analysis (white). VAF: variant allele frequency.

Somatic mutations in *DNMT3A, TET2*, and *ASXL1* (referred to as “DTA” mutations) are commonly found in AML patients but are also seen in clinically asymptomatic individuals with increased prevalence with aging^24-27^. Mutations in these genes are known to not be useful in measuring residual disease in AML^6,12^. Patients with and without DTA mutations were therefore analyzed separately (Fig S5). Overall, 27 patients (60%) had a DTA mutation, and this observation was consistent between the NHLBI relapsed/refractory (median age: 52) and Duke newly diagnosed (median age: 63) AML cohorts. DTA patients expressed significantly lower levels of *WT1* than non-DTA patients (median dCT of 4.8 compared to 3.1 P<0.05) but had greater decrease in *WT1* levels by day 4 (median dCT 7.6 vs. 3.6, P<0.05). 48% of DTA patients achieved CR compared with 33% of non-DTA patients. *NPM1* mutations (n=5) were seen exclusively in DTA patients while *TP53* mutations were seen in both DTA (n=5) and non-DTA (n=4) patients. Only the transcription-related gene class, in non-DTA patients, showed significant difference between day 1 and 4 VAF levels (Figure S5CD).

### Tracking post-treatment MRD in CR patients with NGS

Given the inability of targeted DNA sequencing of blood early during therapy to predict response to a cycle of intensive chemotherapy, we also investigated the utility of this technique in predicting post-remission relapse. Longitudinal blood samples were available from two patients from this cohort both of whom achieved CR. The first patient had no change in the ratios of wild type to mutated sequence of five genes between day 1 and 4 of therapy, despite decreasing WBC count from 60,000/ul to 10,000/ul during this period and subsequently achievement of a durable CR. Mutation levels remained negligible however during a durable remission lasting at least two years (Figure 4A). In the second patient, detectable *KRAS* mutant in blood decreased during the first four days of therapy while the *DNMT3A* mutant remained stable. During remission however both mutations were undetectable, returning at the time of relapse together with the emergence of a second *KRAS* mutation (Figure 4B). DNA sequencing may have greater utility for tracking of AML MRD post-treatment rather than predicting response during therapy.

**Figure 4.**
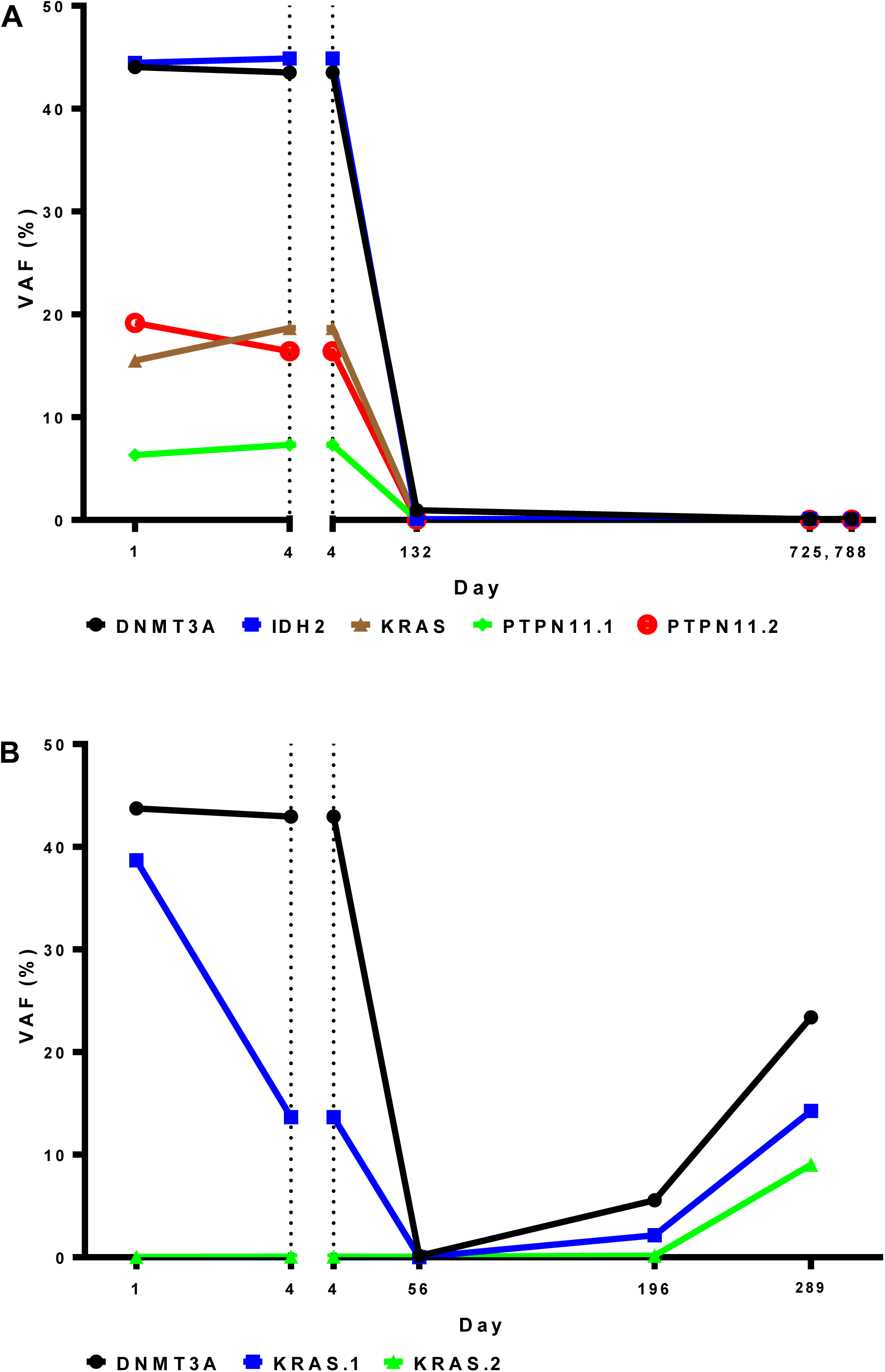
Targeted DNA sequencing for residual disease may be more informative after, rather than during, initial chemotherapy. A DNA sequencing panel customized to patient-specific variants was used to for analysis of longitudinal blood samples from during and after induction therapy in two patients who achieved complete remission. Variant frequency was unchanged in the first patient between day 1 and 4 during therapy, despite achievement of a durable complete remission (A). In the second patient a *DNMT3A* variant changed little during therapy, in contrast to a *KRAS* variant that was markedly reduced. Relapse occurred on day 154. Both mutations were detected In blood from day 196 followed by the emergence of an second *KRAS* mutation on day 289. VAF: Variant allele frequency.

### DNA sequencing from blood versus bone marrow samples

In a subset of 22 patients confirmatory sequencing was also performed on pre-treatment bone marrow samples (10/10 NHLBI, 12/35 Duke). Concordance between the number of variants identified and the VAF of each detected variant in blood compared with bone marrow was assessed (Figure 5A). In the 15 patients with a pre-treatment WBC of at least 2500/ul only a single variant identified from bone marrow was not detected from blood (of 41 variants identified in total, ie: 98%). Notably, there was good correlation in the VAF determined for each variant from both tissue sources in these patients. Conversely, five variants were identified only from blood and not from bone marrow.

**Figure 5.**
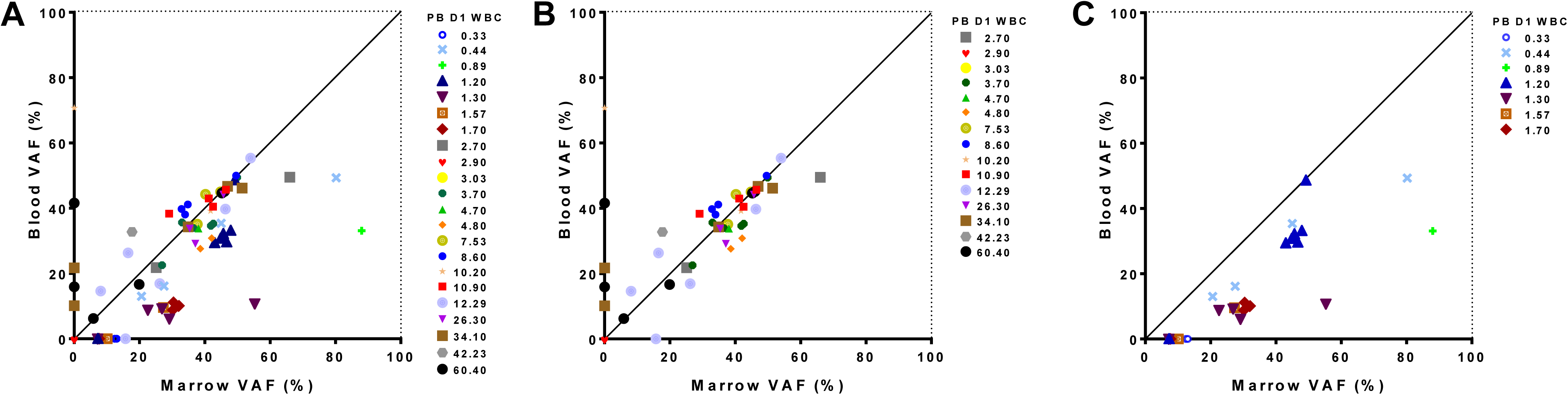
Relationship between sequence variants detected from blood and marrow sample in AML patients. Results of targeted DNA sequencing at day 1 of treatment show good concordance with outliers associated to extremes of white blood cell (WBC) count in blood. Variant allele frequency detected from in blood and marrow is shown for all 22 patients evaluated (A), just those patients with WBC greater than 2500/ul (B) or just those patients with WBC less than 2500/ul in blood (C). Leukopenic patients had lower VAF in blood for variants identified from marrow. One patient had no detectable mutations in either blood or marrow. PB D1 WBC: White blood cell count in blood on Day 1. VAF: Variant allele frequency.

Interestingly, the presence or absence of mutations in the blood versus bone marrow was often correlated with WBC. Two patients with high pre-treatment WBC (34,100/ul and 60,400/ul) accounted for four of the five variants variants observed in the blood but not the bone marrow (Figure 5B). Likewise, for 7 patients with pre-treatment WBC less than 2,500/ul, only 20 out of a total of 27 variants detected in bone marrow were also detected from blood (74%), with the consequence that 5 of 7 leukopenic AML patients had an incomplete mutational characterization by DNA sequencing when using blood alone (Figure 5C). Those variants found in marrow but not identified in blood could however be identified in the raw sequencing data but were filtered out by the 5% VAF threshold for variant calling. For all 20 variants identified in both tissues the VAF was lower in blood than marrow in these leukopenic patients.

## Discussion

As we strive to personalize treatments to cancer patients, both in terms of the genetic basis of their cancer and their response to therapy, there is a great interest in earlier assessments of disease burden and characteristics. AML offers a unique opportunity to study the validity of blood-based assessments of residual tumor, as both the primary site of disease (bone marrow) and blood are repeatedly sampled as part of the clinical standard of care. It is increasingly recognized that blood, except in cases of leukopenia or low circulating blast count, may substitute for marrow examination in some circumstances for morphology, cytogenetics and molecular testing in AML patients^28-30^. We show here that blood, even after three days of intensive cytotoxic chemotherapy, can be used to identify most of the MDS/AML-associated DNA variants detectable by targeted sequencing in pre-treatment bone marrow aspirate, providing that pre-treatment white blood cell count was within or above normal range.

Complete remission is a necessary, but often insufficient, step towards long-term cure in AML. Given that the outcome of patients with relapsed and refractory AML is generally poor^31,32^ there is great interest in early interim assessments of likely response to optimize therapy in AML. While the role of bone marrow examination on day 14 of induction therapy remains unclear^30,33-36^, there has been considerable interest in kinetics of early blast clearance in blood during induction therapy as a prognostic factor^35,37-39^. Persistence of variants detected by targeted sequencing in AML patients in CR after treatment has significant independent prognostic value for both relapse and survival^6^. It was therefore intriguing to consider if such molecular assessments, performed during cytotoxic therapy and prior to any response assessment, could offer a “real-time” evaluation of treatment efficacy and extremely early identification of treatment failure. We show that molecular testing of blood on treatment day 4, by either qPCr assessment of *WT1* expression or DNA sequencing for common MDS/AML variants, is not predictive of clinical response to intensive cytotoxic therapy in AML. Surprisingly, all variants found by DNA sequencing in blood on day one remained detectable on the fourth day of intensive chemotherapy. Remarkably the ratio of mutated to wild-type sequence was often maintained during this therapy despite considerable reductions in white blood cell count. This finding of stability was not limited to potential germline mutants or to variants in *DNMT3A, TET2* and *ASXL1*. The observed stability in VAF during cytotoxic therapy may suggest a limited therapeutic index for clones circulating in blood containing these mutations, although similar studies in patients receiving highly effective and specific therapy would be needed to prove this definitively. The kinetics of *NPM1* and *TP53* variant burden early during chemotherapy however did appear to exhibit consistent trends during therapy in this cohort and are markers potentially worthy of future study.

This study has several limitations. Firstly, we did not perform germline sequencing to allow categorization of identified variants by DNA sequencing as somatic. This was intentional, given the translational nature of this study, we wished to replicate testing as commonly performed in clinical practice. Importantly germline mutations would not be expected to change during chemotherapy so they would not be informative for, or contribute to, variant kinetic analysis. Additionally, the majority of variants identified had a VAF of less than 40%, and 42 of 43 assessed patients had at least one variant with VAF less than 40% detectable in blood at baseline (Figure S4). Secondly, AML represents a wide range of myeloid malignancies with many patterns of genetic etiology whereas we used a targeted sequencing panel designed to detect just 49 commonly mutated genes or gene regions in MDS/AML (Table S3). While more comprehensive approaches have been used to characterize potential genetic variants associated with the leukemic clone that may be detectable before and after therapy^11^, we felt assessment of the most recurrently observed mutated regions was most easily translatable. Finally, we demonstrate consistent observations in two independent cohorts, at two different stages of disease treated with different intensive chemotherapy regimens. Given the genetic heterogeneity of this disease however it is possible that larger cohorts of patients would identify additional trends and will certainly be needed to quantify any benefit associated with tracking the two targets, *TP53* and *NPM1*, we identify as potential candidates for monitoring during intensive therapy.

In conclusion, our study demonstrates that molecular testing of peripheral blood during the first three days of AML intensive chemotherapy does not appear to be predictive of clinical response. Indeed, we show that the majority of variants identified prior to treatment are still present and often at similar ratios of mutant to wild-type often despite considerable cytotoxic effect of therapy. Validation in a larger cohort is needed to confirm the utility of monitoring *NPM1* and *TP53* variant kinetics in blood early during AML treatment in addition to their current use as pre-treatment predictive markers. Consistent with reports using other modalities we show that blood may substitute for bone marrow for targeted DNA sequencing in AML patients, although this approach may be suboptimal in those with leukopenia pre-treatment. Longitudinal assessment of molecular MRD during follow-up time points after completion of initial cytotoxic induction therapy may have greater clinical utility than evaluation of very early time points during AML treatment.

## Acknowledgments

This work was supported by the Intramural Research Program of the National Heart, Lung, and Blood Institute of, and grant 5KL2TR001115-03 from, the National Institutes of Health. The authors wish to acknowledge the assistance of Dr. Yuesheng Li and his team at the NHLBI Sequencing and Genomics Core, Dr. Yanqin Yang for bioinformatics support, Elena Cho, Therese Intrater, Sophie Grasmeder, Tat’Yana Worthy and Debbie Draper for research nurse support, Dr. Qingguo Liu, Lemlem Alemu and Aasheen Qadri for laboratory technical support and to thank the clinical teams, regulatory staff and patients at NIH and Duke.

## References

1. Dohner H, Estey E, Grimwade D, et al: Diagnosis and management of AML in adults: 2017 ELN recommendations from an international expert panel. Blood 129:424–447, 2017

2. Schuurhuis GJ, Heuser M, Freeman S, et al: Minimal/measurable residual disease in AML: a consensus document from the European LeukemiaNet MRD Working Party. Blood 131:1275–1291, 2018

3. Arber DA, Borowitz MJ, Cessna M, et al: Initial Diagnostic Workup of Acute Leukemia: Guideline From the College of American Pathologists and the American Society of Hematology. Arch Pathol Lab Med 141:1342–1393, 2017

4. Hourigan CS, Karp JE: Minimal residual disease in acute myeloid leukaemia. Nat Rev Clin Oncol 10:460–71, 2013

5. Zhou Y, Othus M, Walter RB, et al: Deep NPM1 Sequencing Following Allogeneic Hematopoietic Cell Transplantation Improves Risk Assessment in Adults with NPM1-Mutated AML. Biol Blood Marrow Transplant, 2018

6. Jongen-Lavrencic M, Grob T, Hanekamp D, et al: Molecular Minimal Residual Disease in Acute Myeloid Leukemia. N Engl J Med 378:1189–1199, 2018

7. Malmberg EB, Stahlman S, Rehammar A, et al: Patient-tailored analysis of minimal residual disease in acute myeloid leukemia using next-generation sequencing. Eur J Haematol 98:26–37, 2017

8. Gaksch L, Kashofer K, Heitzer E, et al: Residual disease detection using targeted parallel sequencing predicts relapse in cytogenetically normal acute myeloid leukemia. Am J Hematol 93:23–30, 2018

9. Morita K, Kantarjian HM, Wang F, et al: Clearance of Somatic Mutations at Remission and the Risk of Relapse in Acute Myeloid Leukemia. J Clin Oncol 36:1788–1797, 2018

10. Rothenberg-Thurley M, Amler S, Goerlich D, et al: Persistence of pre-leukemic clones during first remission and risk of relapse in acute myeloid leukemia. Leukemia 32:1598–1608, 2018

11. Klco JM, Miller CA, Griffith M, et al: Association Between Mutation Clearance After Induction Therapy and Outcomes in Acute Myeloid Leukemia. Jama 314:811–22, 2015

12. Debarri H, Lebon D, Roumier C, et al: IDH1/2 but not DNMT3A mutations are suitable targets for minimal residual disease monitoring in acute myeloid leukemia patients: a study by the Acute Leukemia French Association. Oncotarget 6:42345–53, 2015

13. Hourigan CS, Gale RP, Gormley NJ, et al: Measurable residual disease testing in acute myeloid leukaemia. Leukemia 31:1482–1490, 2017

14. Terwijn M, van Putten WL, Kelder A, et al: High prognostic impact of flow cytometric minimal residual disease detection in acute myeloid leukemia: data from the HOVON/SAKK AML 42A study. J Clin Oncol 31:3889–97, 2013

15. Ivey A, Hills RK, Simpson MA, et al: Assessment of Minimal Residual Disease in Standard-Risk AML. N Engl J Med 374:422–33, 2016

16. Freeman SD, Hills RK, Virgo P, et al: Measurable Residual Disease at Induction Redefines Partial Response in Acute Myeloid Leukemia and Stratifies Outcomes in Patients at Standard Risk Without NPM1 Mutations. J Clin Oncol 36:1486–1497, 2018

17. Buckley SA, Wood BL, Othus M, et al: Minimal residual disease prior to allogeneic hematopoietic cell transplantation in acute myeloid leukemia: a meta-analysis. Haematologica 102:865–873, 2017

18. Araki D, Wood BL, Othus M, et al: Allogeneic Hematopoietic Cell Transplantation for Acute Myeloid Leukemia: Time to Move Toward a Minimal Residual Disease-Based Definition of Complete Remission? J Clin Oncol 34:329–36, 2016

19. Hourigan CS, Goswami M, Battiwalla M, et al: When the Minimal Becomes Measurable. J Clin Oncol 34:2557–8, 2016

20. Cilloni D, Renneville A, Hermitte F, et al: Real-time quantitative polymerase chain reaction detection of minimal residual disease by standardized WT1 assay to enhance risk stratification in acute myeloid leukemia: a European LeukemiaNet study. J Clin Oncol 27:5195–201, 2009

21. Goswami M, Hensel N, Smith BD, et al: Expression of putative targets of immunotherapy in acute myeloid leukemia and healthy tissues. Leukemia 28:1167–70, 2014

22. Ley TJ, Miller C, Ding L, et al: Genomic and epigenomic landscapes of adult de novo acute myeloid leukemia. N Engl J Med 368:2059–74, 2013

23. Papaemmanuil E, Gerstung M, Bullinger L, et al: Genomic Classification and Prognosis in Acute Myeloid Leukemia. N Engl J Med 374:2209–2221, 2016

24. Genovese G, Kahler AK, Handsaker RE, et al: Clonal hematopoiesis and blood-cancer risk inferred from blood DNA sequence. N Engl J Med 371:2477–87, 2014

25. Zink F, Stacey SN, Norddahl GL, et al: Clonal hematopoiesis, with and without candidate driver mutations, is common in the elderly. Blood 130:742–752, 2017

26. Shlush LI: Age-related clonal hematopoiesis. Blood 131:496–504, 2018

27. Jaiswal S, Fontanillas P, Flannick J, et al: Age-related clonal hematopoiesis associated with adverse outcomes. N Engl J Med 371:2488–98, 2014

28. Weinkauff R, Estey EH, Starostik P, et al: Use of peripheral blood blasts vs bone marrow blasts for diagnosis of acute leukemia. Am J Clin Pathol 111:733–40, 1999

29. Tong WG, Sandhu VK, Wood BL, et al: Correlation between peripheral blood and bone marrow regarding FLT3-ITD and NPM1 mutational status in patients with acute myeloid leukemia. Haematologica 100:e97–8, 2015

30. Percival ME, Lai C, Estey E, et al: Bone marrow evaluation for diagnosis and monitoring of acute myeloid leukemia. Blood Rev 31:185–192, 2017

31. Breems DA, Van Putten WL, Huijgens PC, et al: Prognostic index for adult patients with acute myeloid leukemia in first relapse. J Clin Oncol 23:1969–78, 2005

32. Ramos NR, Mo CC, Karp JE, et al: Current Approaches in the Treatment of Relapsed and Refractory Acute Myeloid Leukemia. J Clin Med 4:665–95, 2015

33. Morris TA, DeCastro CM, Diehl LF, et al: Re-induction therapy decisions based on day 14 bone marrow biopsy in acute myeloid leukemia. Leuk Res 37:28–31, 2013

34. Hussein K, Jahagirdar B, Gupta P, et al: Day 14 bone marrow biopsy in predicting complete remission and survival in acute myeloid leukemia. Am J Hematol 83:446–50, 2008

35. Yanada M, Borthakur G, Ravandi F, et al: Kinetics of bone marrow blasts during induction and achievement of complete remission in acute myeloid leukemia. Haematologica 93:1263–5, 2008

36. Yezefski T, Xie H, Walter R, et al: Value of routine ‘day 14’ marrow exam in newly diagnosed AML. Leukemia 29:247–9, 2015

37. Kern W, Haferlach T, Schoch C, et al: Early blast clearance by remission induction therapy is a major independent prognostic factor for both achievement of complete remission and long-term outcome in acute myeloid leukemia: data from the German AML Cooperative Group (AMLCG) 1992 Trial. Blood 101:64–70, 2003

38. Gianfaldoni G, Mannelli F, Baccini M, et al: Clearance of leukaemic blasts from peripheral blood during standard induction treatment predicts the bone marrow response in acute myeloid leukaemia: a pilot study. Br J Haematol 134:54–7, 2006

39. Vainstein V, Buckley SA, Shukron O, et al: Rapid rate of peripheral blood blast clearance accurately predicts complete remission in acute myeloid leukemia. Leukemia 28:713–6, 2014

40. Goswami M, McGowan KS, Lu K, et al: A multigene array for measurable residual disease detection in AML patients undergoing SCT. Bone Marrow Transplant 50:642–51, 2015

